# Protoplast fusion in *Bacillus* species produces frequent, unbiased, genome-wide homologous recombination

**DOI:** 10.1101/2020.10.06.328625

**Authors:** Leah H. Burdick, Jared C. Streich, Delyana P. Vasileva, Dawn M. Klingeman, Hari B. Chhetri, J. Christopher Ellis, Dan M. Close, Daniel A. Jacobson, Joshua K. Michener

## Abstract

In eukaryotes, fine-scale maps of meiotic recombination events have greatly advanced our understanding of the factors that affect genomic variation patterns and evolution of traits. However, in bacteria that lack natural systems for sexual reproduction, unbiased characterization of recombination landscapes has remained challenging due to variable rates of genetic exchange and influence of natural selection. Here, to overcome these limitations and to gain a genome-wide view on recombination, we crossed *Bacillus* strains with different genetic distances using protoplast fusion. The offspring displayed complex inheritance patterns with one of the parents consistently contributing the major part of the chromosome backbone and multiple unselected fragments originating from the second parent. Computational analyses suggested that this bias is due to the action of restriction-modification systems whereas genome features like GC content and local nucleotide identity did not affect distribution of recombination events around the chromosome. Furthermore, we found that the intensity of recombination is uniform across the genome without concentration into hotspots. Unexpectedly, our results revealed that large species-level genetic distance did not affect key recombination parameters. Our study provides a new insight into the dynamics of recombination in bacteria and a platform for studying recombination patterns in diverse bacterial species.

## INTRODUCTION

Homologous recombination in the form of uptake and integration of DNA from exogenous sources has played a profound role in shaping microbial evolution and speciation (1). However, genetic transfer and recombination are rare in natural bacterial populations and thus difficult to characterize in detail. While a number of computational methods have been developed to estimate the relative rates and distribution of recombination events based on genome sequences of extant bacteria (2), these analyses are confounded by historical selection on recombinant strains. Direct measurements of recombination parameters on a genome-wide scale are technically challenging because recombination patterns can be significantly affected by efficiencies and mechanistic specificities of DNA transfer. To date, most experimental estimates of recombination rates have been conducted by *in vitro* transformation of naturally competent bacteria (3–5), but under these conditions transfer is typically limited to only small regions of the genome. A greater portion of chromosomal DNA spanning hundreds of genes could be exchanged between bacteria through some unconventional conjugal mechanisms resembling Hfr-based transfer in *Escherichia coli* (6–10). For example, mycobacterial distributive conjugal transfer and mycoplasma chromosomal transfer can promote simultaneous transfer of multiple large donor chromosomal fragments to the recipient cells, creating chimeric transconjugant genomes with unique recombination landscapes. Although these studies have provided an invaluable insight into the genetics of recombination, they were restricted in scale and scope due to computational limitations. Genome-level perspective on recombination in bacteria is still lacking and, as a result, we do not fully understand how features of the genomic environment affect recombination rates.

High frequencies of genetic transfer and recombination on a genome-wide scale in bacteria can be achieved by protoplast fusion (11). In this classical genetic engineering method, bacterial cells are stripped of their outer layer and chemically fused together, allowing recombination between the parental chromosomes. Originally used for routine genetic manipulation, protoplast fusion has been widely adopted as a strategy to generate microorganisms with improved phenotypes for biotechnological applications by combining beneficial alleles from different strains and even species (12, 13). For instance, combinatorial shuffling of complete genomes by recursive fusion of protoplast populations has been employed to engineer multigenic traits for which the underlying molecular mechanisms are poorly understood, such as tolerance to stress conditions and production of diverse metabolites (14–16). Multiple crossover events are generally assumed to occur across the entire genome during this process, giving rise to mosaic chromosomes with unique phenotypic potential, analogous to meiotic recombination products in sexually reproducing organisms. Surprisingly, the exact nature of the chromosomal rearrangements resulting from large-scale shuffling experiments has received little attention and to date there are few studies reporting detailed analyses of sequenced bacterial shuffled genomes (17–19). Furthermore, due to strong selective pressure for the desired phenotypes, these analyses could not capture the full extent of recombination occurring between the parental chromosomes in protoplast fusants.

Mosaic genomes generated by DNA shuffling provide a unique source to investigate the genomics of recombination. In this work, we generated recombinant progeny from protoplast fusion between pairs of *Bacillus* strains with varying degrees of nucleotide identity. We built fine-scale recombination maps using next-generation sequencing and developed a computational pipeline to gain a deeper insight into how genomic sequence parameters affect dynamics of recombination events. Our results revealed that protoplast fusion generates multiple recombination events distributed across the genome with bias towards one of the parents and no other regional biases. Strikingly, our data showed that core features of homologous recombination were unaffected by large differences in nucleotide identity between parental strains. This work might aid in a better understanding of bacterial evolution in natural systems as well as provide potential insights into the use of genome shuffling for improving cellular function.

## MATERIAL AND METHODS

### Strains and chemicals

*Bacillus* strains used in this work are summarized in Table 1. Parental cell lines were initially grown in low-salt LB medium (casein digest peptone 10 g/l, NaCl 5 g/l, yeast extract 5 g/l) with antibiotic selection as appropriate. All cells were grown at 37°C with liquid cultures kept at 250 rpm rotation. Antibiotics used include kanamycin sulfate (50 μg/ml), and erythromycin (20 μg/ml) used in concert with lincomycin (12.5 μg/ml). During the shuffling procedure, cells were washed and maintained in SMM buffer (20) consisting of 0.5M sucrose, 20 mM MgCl_2_, and 20 mM maleic acid. PEG buffer to induce protoplast fusion consisted of SMM buffer supplemented with 35% PEG 6,000 and 10 mM CaCl_2_ (21). Newly shuffled cells were plated on DM3 recovery medium (22) and subsequently plated on minimal medium (MM) (23) or LB agar plates. Media were supplemented with antibiotics as described above and with tryptophan (400 μM), histidine (300 μM), and methionine (1 mM) as needed for various auxotrophic strains.

**Table 1.** Strains used in this study.

| Strain | Genotype | Phenotype | References |
| --- | --- | --- | --- |
| *B. subtilis* subsp. *subtilis* 168 | *trpC2* | *trp*<sup>−</sup> | (51–53) |
| *B. subtilis* subsp. *subtilis* RO-NN-1 | Wild-type | Wild-type | (54, 55) |
| *B. subtilis* subsp. *subtilis* NCIB3610 | Wild-type | Wild-type | (52, 56) |
| *B. subtilis* subsp. *spizizenii* TU-B-10 | Wild-type | Wild-type | (54, 57) |
| *B. mojavensis* RO-H-1 | Wild-type | Wild-type | (54, 55) |
| *B. velezensis* FZB42 | Wild-type | Wild-type | (58, 59) |
| BKE13180 | 168 $\Delta metE::erm$ | *trp<sup>-</sup> met<sup>-</sup> erm<sup>R</sup>* | (24) |
| BKK34900 | 168 $\Delta hisB::kan$ | *trp<sup>-</sup> his<sup>-</sup> kan<sup>R</sup>* | (24) |
| JMB1 | RO-NN-1 $\Delta hisB::kan$ | *his<sup>-</sup> kan<sup>R</sup>* | This work |
| JMB3 | RO-NN-1 $\Delta metE::erm$ | *met<sup>-</sup> erm<sup>R</sup>* | This work |
| JMB60 | RO-NN-1 $\Delta metE::erm$<br>$\Delta hisB::kan$ | *his<sup>-</sup> met<sup>-</sup> kan<sup>R</sup> erm<sup>R</sup>* | This work |
| JMB12-JMB29<br>(Figure 1B/2B, blue) | 168 $\Delta metE::erm$ x<br>RO-NN-1 $\Delta hisB::kan$ | Wild-type | This work |
| JMB6-JMB11<br>JMBP3A1-JMBP3A12<br>(Figure 1C/2B, orange) | 168 $\Delta metE::erm$ x<br>RO-NN-1 $\Delta hisB::kan$ | *his<sup>-</sup> met<sup>-</sup> kan<sup>R</sup> erm<sup>R</sup>* | This work |
| JMBP3C4-JMBP3D5<br>(Figure 1D/2B, pink) | 168 $\Delta hisB::kan$ x<br>RO-NN-1 $\Delta metE::erm$ | *his<sup>-</sup> met<sup>-</sup> kan<sup>R</sup> erm<sup>R</sup>* | This work |
| JMBP4D1-JMBP4E4<br>(Figure 4, green) | RO-NN-1 $\Delta metE::erm$<br>$\Delta hisB::kan$ x<br>NCIB3610 | *his<sup>-</sup> kan<sup>R</sup>* | This work |
| JMBP4E5-JMBP4F8<br>(Figure 4, yellow) | RO-NN-1 $\Delta metE::erm$<br>$\Delta hisB::kan$ x<br>NCIB3610 | *met<sup>-</sup> erm<sup>R</sup>* | This work |
| JMB61-JMB75<br>(Figure 4, red) | RO-NN-1 $\Delta metE::erm$<br>$\Delta hisB::kan$ x TU-B-10 | *his<sup>-</sup> kan<sup>R</sup>* | This work |
| JMB76-JMB89<br>(Figure 4, blue) | RO-NN-1 $\Delta metE::erm$<br>$\Delta hisB::kan$ x TU-B-10 | *met<sup>-</sup> erm<sup>R</sup>* | This work |
| JMBP3D8-JMBP3E12<br>(Figure 4, orange) | RO-NN-1 $\Delta metE::erm$<br>$\Delta hisB::kan$ x RO-H-1 | *his<sup>-</sup> kan<sup>R</sup>* | This work |
| JMBP3F1-JMBP3G6<br>(Figure 4, pink) | RO-NN-1 $\Delta metE::erm$<br>$\Delta hisB::kan$ x RO-H-1 | *met<sup>-</sup> erm<sup>R</sup>* | This work |

### Strain construction

Strains BKK34900 (168 Δ*hisB*::*kan*) and BKE13180 (168 Δ*metE*::*erm*) were provided by the BGSC (24). The allele replacement constructs were amplified from genomic DNA of these strains using primers hisB-FL: 5’-CAATTGCCGGATATAATGTAAAAGCAC-3’ /hisB-RL: 5’-ATATGATTGCCGGACCGAGTG AAATC-3’ and metE-FL: 5’-CATGCCTGATCCTTTTAATATTCTTTCTTATTG-3’ /metE-RL: 5’-GCTATG AAGAAGAATCATTTCAAAGAAAG-3’, respectively. Strain RO-NN-1 was then transformed with these PCR products by natural competence, following standard protocols (24). Transformed strains were selected using LB medium containing the appropriate antibiotic and verified by colony PCR. Mutant strains were then resequenced as described below.

A double mutant strain of RO-NN-1, containing both Δ*hisB*::*kan* and Δ*metE*::*erm*, was constructed by genome shuffling as described below. This strain was then verified by whole-genome resequencing.

### Genome shuffling

Cells for genome shuffling were grown in selective liquid media overnight, then diluted 100-fold the following morning. Once cultures reached an OD_600_ between 0.4 and 0.6, 5 ml were pelleted by centrifugation for 5 min at 8,000 x g and 25°C and washed three times in 1 ml SMM buffer. DNase I (5 μg/ml) was added to the SMM buffer after initial wash steps. Protoplast formation was accomplished by resuspending washed cells in 1 ml SMM buffer with 1 mg/ml lysozyme followed by incubation at 37°C for one hour. 500 μl of each parental cell line were mixed together after protoplasting and centrifuged for 20 min at 2,000 x g at 12°C. These mixed pools were washed once in SMM buffer, resuspended in PEG buffer, and incubated at room temperature for 20 minutes. Cells were again washed in SMM buffer and resuspended in 100 μl SMM buffer with 1% BSA added. Cells were then plated on DM3 regeneration media and incubated overnight at 37°C. The following day, cells were scraped from regeneration plates and plated to selective media for single colony isolation.

### Strain isolation and sequencing

Individual strains were isolated either by plating serial dilutions or streaking to individual colonies on selective media. Single colonies were then picked and re-streaked to selective plates before being grown to saturation in selective liquid media. Genomic DNA was isolated using the Qiagen DNeasy Blood and Tissue Kit (Qiagen, Valencia, CA) according to the manufacturer’s instructions. DNA for PacBio sequencing was isolated using the same method, but multiple samples were combined and concentrated to obtain higher concentrations. To achieve this, one tenth combined sample volume of 3M sodium acetate was added to pooled DNA, followed by 2.5 volumes of 100% ethanol. This was mixed and incubated at −80°C for 30 minutes. Precipitated DNA was then pelleted by centrifugation at 14,000 rpm for 20 minutes at 4°C, washed with 70% ethanol, and allowed to air dry. DNA was then resuspended in 1/10 TE buffer and stored at −20°C until being shipped on dry ice to the University of Maryland for PacBio sequencing.

For strain resequencing, Nextera XT libraries (Illumina, San Diego, CA) were generated from purified DNA of isolated strains according to the manufacturer’s protocol (15031942 v03), stopping after library validation. Final libraries were validated on an Agilent Bioanalyzer (Agilent, Santa Clara, CA) using a DNA7500 chip and concentration was determined on an Invitrogen Qubit (Waltham, MA) with the broad range double stranded DNA assay. Barcoded libraries were pooled and prepared for sequencing following the manufacturer’s recommended protocol (15039740v09, Standard Normalization). One paired end sequencing run (2 x 301) was completed on an Illumina MiSeq instrument (Illumina, San Diego, CA) using v3 chemistry. Illumina resequencing of strains generated by crossing RO-NN-1 Δ*metE*::*erm* Δ*hisB*::*kan* with wild-type isolates was performed commercially (SNPSaurus, Eugene, OR). PacBio sequencing was performed by the University of Maryland Institute for Genome Sciences (Baltimore, MD).

### Variant Calling

Fastq files from sequencing were first processed with Trimmomatic for phred base-pair quality. Reads that lost a paired read from phred filtering were removed. Reads that were shorter than 38 base-pairs were removed to reduce the quantity of non-uniquely mapping reads. Individuals were independently run through a variant calling pipeline using software current at the time the project started: BWA v0.7.17, Samtools v1.8, Picard v2.20.8, GATK v3.8.0, VCFTools v0.1.15, BCFTools v1.9, PLINK v1.9.0, and in-house R scripts (25–30). Reads were aligned through BWA MEM to generate .sam files (Sam files). Samtools was then used to create compressed .bam files (Bam files) for further processing. Bam files were then parsed by samtools for uniquely mapping reads to a single locus, while multi-loci mapping reads were removed. Samtools was next used to order reads by their individual genome mapping coordinate and their read groups replaced. After removing non-mappable reads, and remaining reads ordered and properly annotated, Bam files were scanned for duplicate calls with Samtools and then were indexed via Picard. Polished Bam file reads were run through GATK HaplotypeCaller as haploids with “-ploidy 1”. BCFTools was used to filter low coverage variants, requiring a minimum read depth of 12 to confirm the variant. GATK’s HaplotypeCaller function will only annotate the most common variant in haploid organisms, and since sequencing errors are rare, only variants with several reads (≥20) are marked in VCF files. Variants were also BCFTool filtered for a genotype quality of p<0.1×10^−6^ to ensure the chance of a false variant was less than 1:100,000. A random subset of individuals were then scanned by eye to check for variants in low coverage areas, that no low-coverage variants were marked, and no biallelic states were present. Final bioinformatic analysis was done in R v3.5.0 using Plink ped/map file format.

### Genomic Feature Analysis

#### Parent Detection and Filtering

Within each shuffled population variants were first called against each parent reference genome. However, in every recombinant population the parent strain RO-NN-1 remained the dominant contributor to offspring genomes, and thus was used for all further variant calling and genome analysis. In each population variants were encoded as “0” for RO-NN-1 and “2 as the recessive parent. Variants called in both parents at a single position are likely sequencing errors that arose during laboratory processing or DNA sequencing. Markers not present in at least one offspring were also removed. Any variant found in one parent and one individual were kept for recombination and insertion analysis methods. Lists for differential variants between parents were used for permutation testing (described below).

#### Insert Size and Frequency

Post variant encoding, insert size was calculated based on the number of base pairs between continuous variants from the recessive parent. The positions and lengths of insertions from recessive parents were calculated for each individual by counting continuous strings of non-reference variants and their distance in bases along the genome. Features of shuffled genomes were visualized by the R package ‘BioCircos’ and standard plotting libraries (31). The quantity of insertions per strain was similarly quantified by totalling each individual’s strings of recessive markers. The distributions were tested for normality using the Shapiro-Wilk tests. The means and standard deviations were compared with T-tests, F-tests, Wilcoxon-Tests, and Kolmogorov-Smirnov tests for significance.

#### Population Level Genome Feature Analysis

Read mapping statistics were calculated using VCFTools and the RO-NN-1 reference genome as the dominant parent (26). Read depth per shuffle was calculated using ‘--depth’ for population level read depth. Likewise, VCFTools function “--mean-depth” was used for broad read depth, and ‘--site-depth’ was used for variant sequence depth per individual in each shuffle. Site mean depth was calculated by VCFTools ‘--site-mean-depth’ function to obtain per sample mean sequence depth.

#### Permutation Testing Against Genome Features

Recombined positions in the genome were examined against other extractable genomic features. IGV v2.3.5 was used for genomic feature extraction of a known methylation motif (GAYGNNNNNNCTT) and GC content (32). Additionally, known gene positions within the dominant reference parent RO-NN-1 were also used in testing variants involved in insertion detection. In each permutation test, a two-stage random number generator was used; the first seed number to create a list of random numbers, that was then used to create a second set of random numbers each used once in a single iteration within tests. In each test, positions of genomic features were compared to randomly generated lists of genomic positions to test if insertions between parents have statistical significance to single nucleotide polymorphisms (SNPs)/variants, methylation motifs, or GC content. Each test was run against 1,000 randomly generated subsets to create a p-value significance level of 0.0001. Iteration subsets of random test positions were based on the number features detected. For instance, 1,066 methylation motifs exist in the RO-NN-1 genome, thus per each iteration 1,066 random positions were used.

#### Methylation to Insertion Testing

To investigate if methylation sites are closer than random to insertion sites we compared “*distance in base pairs from methylation motifs to random positions*” to “*base pair distance of motifs to insertion sites*”. A list of randomly generated genome positions was created to draw subsets per iteration equal to the number of insertion events per population. In each iteration the distance to a methylation was calculated to a randomly drawn genome position to create a distribution of randomly drawn base pair lengths. Then, subsets of the 1,066 known methylation motifs were drawn per iteration and base pair distance was calculated to the nearest 5’ or 3’ end of an insertion event (see Supplementary Figure S1).

#### Insertion Events to Random Position Testing

Insertion events could be biased toward specific positions within the genome. To test this, we generated two lists of random genome positions and calculated base pair distance between pairs of positions. For each random position in data set one, we determined the distance to the closest randomly drawn position in the second random set. Then we randomly drew positions in the genome and calculated distance to the nearest 5’ or 3’ insertion event.

#### GC Content Permutation Testing

Approximately 46% of the *B. subtilis* genome is G or C, thus proximity in bases to the nearest G or C is not meaningful. Two similar tests for GC content correlation to insertion positions were implemented. One test examined uni-directional outward GC content away from insertion sites; from the 5’ insertion then examining increasing windows beforehand (3’ to 5’), and the 3’ end of the insertion expanding forward (5’ to 3’). GC content was measured by percent GC at increasing increments through exponentially increasing windows of 2^n^ bases, *n* = 2:12 (2^2^ from 2^12^; 4 bases to 4,096 bases). The same test was performed on randomly generated insertions, unique to each iteration, and the percent GC was calculated using the same exponential scan pattern as variants. To generate a list of random insertions with comparable insertion lengths, random markers were chosen from a list of known variant sites between the two parents as the 5’ end. To get a comparable 3’ marker as the insertion switch point, actual insertion sizes were randomly drawn and assigned to 5’ variants and the closest 3’ differential variant was chosen in either direction, thus creating the most similar possible insertion size to an observed insertion size. To generate *in-silico* variants required the use of the R package “ecodist” (33). A very similar test was performed scanning GC content, but in both 5’ and 3’ directions from insertion ends (scanning away and into the insertion markers). A smaller set of windows was used since the chance of double counting GC content exists within the boundaries of *in-silico* simulated insertions. When building simulated insertions, insertion sizes that were smaller than 1024 were removed. Thus, window sizes considered ranged from 4^n^, *n* = 1:4 (4 bases to 256 bases). Limiting the GC content scan within insertion sites to 256 bases means that up to 50% of the insertion site was scanned for %GC content.

#### Wavelet Analysis for Population Features and a Range of Complexities

Wavelet transforms can analyze signal-based data by expanding 2 dimensional data into 3 dimensional space at varying scales to reveal otherwise cryptic patterns. The underlying theory of wavelet analysis is to overlay an organized specific wave of designated length and area over a signal series to find differences in area annotated as coefficients. Wavelets can find patterns or quantify “how much of a peak” is present at a region of a signal that is not immediately obvious to the human eye, and scanned at varying scales/window sizes of data (34-35). Within this study we implemented a Continuous Wavelet Transform (CWT) using the Ricker Wavelet as the mother wavelet to identify regions of the genome with differing characteristics of recombinant loci and potential hot and cold locations across recombinant populations. Ricker wavelets are ideal for this scan type since they target one specific location relative only to immediate up and downstream signal, being they are composed of three parts with a total area of zero (two negative peaks with area = −0.5 flanking a single positive peak with area = 1). Below is the wavelet transform that returns the wavelet coefficients *W(s,τ)* that are calculated across scales (s) and translation along the genome as *τ* (shifts across the x-axis) (36).

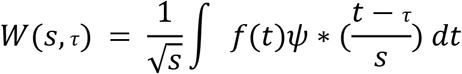

The resulting coefficients will indicate at specific scales the *quantity of peak* present. Wavelet analysis was performed using the R statistical programming language 3.5.0 and the ‘wmtsa’ package was used for wavelet transform analysis (37-38). Genomic data was encoded as “0” and “2” and only for relevant positions such binary transition states collapsed only to varying positions doesn’t lend well to signal processing, thus variant data was modified in two ways (35). First, all variant positions were summed across the population to a single vector and spread out to their actual position, where absence of a variant was annotated as a zero. Secondly, data was binned down to approximately 4,010 data points (100x reduction) depending on the genome marker positions of each population. Once the data was transformed to amenable wavelet analysis qualities the locations with differing areas to the mean with either higher or lower than expected values were revealed.

## RESULTS AND DISCUSSION

### Genomic consequences of genome shuffling

Successful genome shuffling is typically assessed through simultaneous selection for markers present in both parents. To make this strategy more flexible, we replaced biosynthetic genes that are essential for growth in minimal media with antibiotic resistance markers (Figure 1A). This approach allows selection for any of four potential allele combinations. We chose *hisB* and *metE* as biosynthetic genes, since these gene deletions produce known auxotrophies (24) and the genes are roughly opposite in the genome, separated by 2.2 and 2.0 Mb.

**Figure 1:**
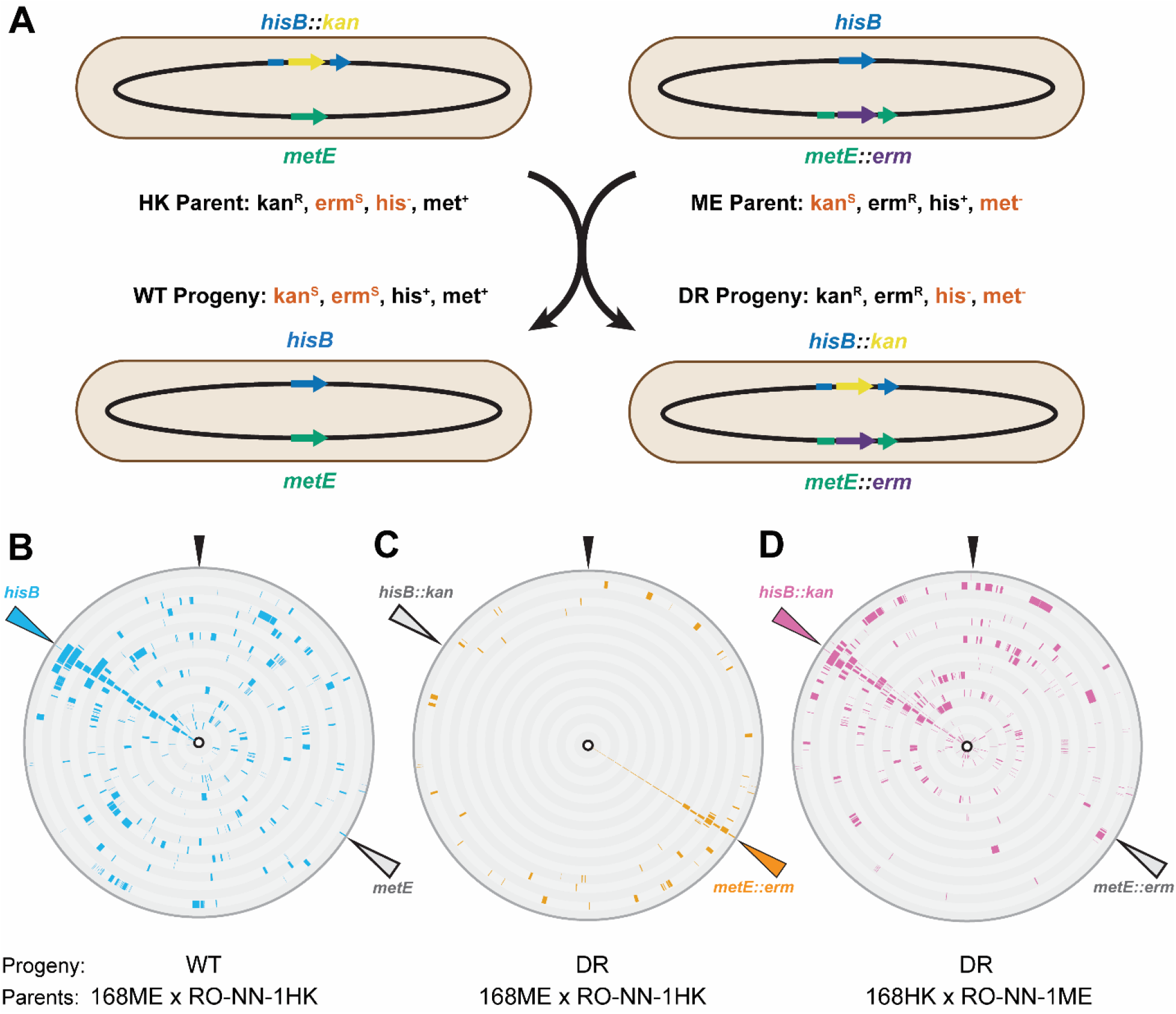
Analysis of genome shuffling in *Bacillus subtilis*. (A) Replacing amino acid biosynthesis genes with antibiotic resistance markers allows flexible identification of recombinant progeny following genome shuffling. (B-D) Crossing mutants of 168 and RO-NN-1 yielded prototrophic (B) and double-resistant (C+D) progeny. Each concentric circle represents a different resequenced individual from this cross. The colored bars indicate sequences recombined from strain 168, with the remaining genomic sequence coming from RO-NN-1. Orange, blue, pink, and grey arrows indicate locations of selection markers. Black arrows indicate the origin of replication. WT: wild-type; DR: double-resistant.

To determine the genome-wide effects of protoplast fusion, we performed reciprocal crosses of 168 Δ*hisB*::*kan* (“168 HK”) x RO-NN-1 Δ*metE*::*erm* (“RO-NN-1 ME”) and RO-NN-1 Δ*hisB*::*kan* (“RO-NN-1 HK”) x 168 Δ*metE*::*erm* (“168 ME”). We then selected recombinant strains containing either both mutant alleles (Δ*hisB*::*kan* Δ*metE*::*erm*, “DR”) or both wild-type alleles (*his*^+^ *met*^+^, “WT”). Eighteen recombinant strains from each combination of shuffle and selection were isolated and resequenced using short-read sequencing. Regrettably, the 168 HK x RO-NN-1 ME prototrophic pool was contaminated by other prototrophic isolates and therefore was not analyzed further. To identify large-scale genome rearrangements, we also sequenced two parental and four recombinant strains using long-read-sequencing. The genome sequences of the recombinant strains were then analyzed computationally to determine the genetic contributions from each parent.

Sequencing results revealed a strong asymmetry in recombination, with one of the parental strains (RO-NN-1 ME or RO-NN-1 HK) contributing the majority of the chromosome of every progeny (Figure 1B-D). All recombinant strains carried the selected marker flanked by different amounts of DNA, ranging from 4 to 124 kb, that originated from the second parent (168 HK or 168 ME). In addition, we detected extensive unselected variation across their genomes with multiple unrelated regions of recombination. Within a single strain, these unselected recombined regions were not distributed evenly around the chromosome, but instead often showed frequent recombination within a relatively small region of the genome (Figure 1B-D and Supplementary Figure S2). Similar inheritance patterns have been observed after conjugation and natural transformation in several bacterial species (3, 4, 6, 10). It has previously been suggested that localized clustering of recombination events might result from uptake of a single large donor DNA fragment followed by multiple rounds of recombination. In this work, however, the entire 168 chromosome was acquired simultaneously, suggesting that frequent local recombination is not an artifact of DNA transport limitations.

Interestingly, genomes of the DR progeny generated by fusion of 168 ME and RO-NN-1 HK protoplasts showed lower levels of complexity compared to the other two progeny populations (Figure 1B-D and Figure 2A). Individual 168 ME x RO-NN-1 HK WT (blue) and 168 HK x RO-NN-1 ME DR (pink) recombinant chromosomes contained a median of 23 and 21 separate 168-derived genome segments, respectively, while 168 ME x RO-NN-1 HK DR (orange) genomes contained a median of 1 fragment originating from 168 ME (Figure 2A). The two sets of 168 ME x RO-NN-1 HK progeny (blue and orange) result from the same fusion and regeneration process, simply plated on different selective media. DR progeny were selected on LB with both antibiotics, while WT progeny were selected on minimal medium. We could not detect significant differences in fitness between any of the parental strains under either growth condition. Thus, the factors that have promoted enrichment of recombinant populations with different levels of heterogeneity remain unclear.

**Figure 2:**
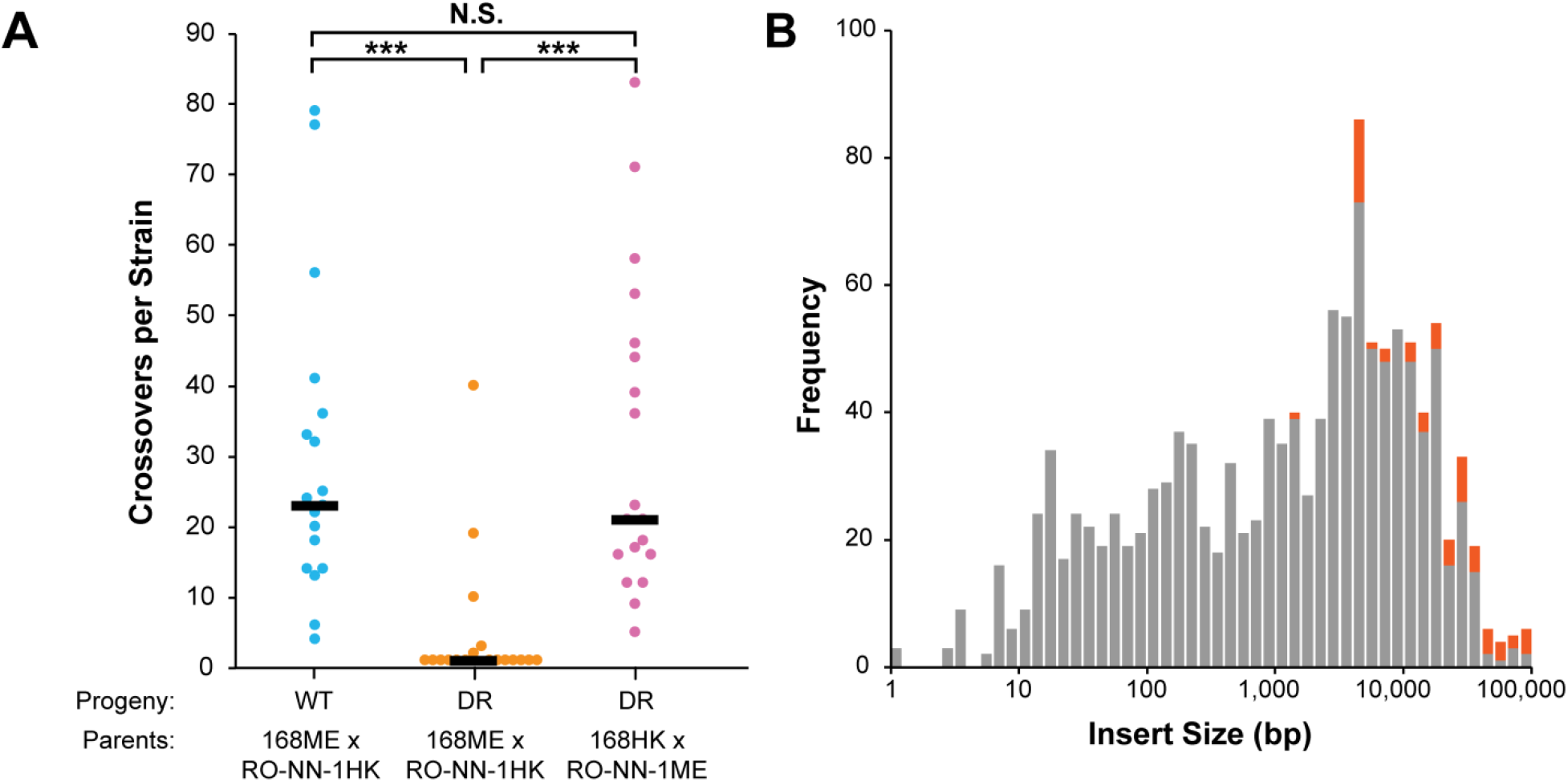
Analysis of recombination frequency and size. (A) The number of recombination events was calculated for each strain in a given pool, representing each strain by a single data point. The median value for each pool is shown with a black line. Shapiro-Wilks tests were used to test distribution normality of number of recombination events showing that each population had non-normally distributed data (Supplementary Table S1). T-tests, F-tests, Kolmogorov-Smirnov and Wilcoxon tests were used to compare variances and distribution means between the three populations (Supplementary Table S2). (B) The distribution of recombination fragment lengths is shown for all three populations combined. Fragments containing the selection marker are indicated in red. WT: wild-type; DR: double-resistant. ***: p<0.001; N.S.: not significant.

Recombination fragment sizes in the three progeny populations were broadly distributed, ranging from a single nucleotide to over 60 kb (Figure 2B). On average, fragments from strain 168 replaced 4.5% (168 ME x RO-NN-1 HK WT), 0.6% (168 ME x RO-NN-1 HK DR), and 3.0% (168 ME x RO-NN-1 HK DR) of the chromosome of RO-NN-1 (Supplementary Table S1). Long-read sequencing of four progeny strains did not identify any large chromosomal rearrangements. Selection ensured that one marker from the 168 parent was necessarily present in the recombinant progeny, and this marker was often integrated as part of a >4 kb segment. The high frequency of recombinant segments of ~4 kb (Figure 2B, red bars) is partly due to this bias.

Recombination can only be detected when it results in a genetic change. The chromosomes of strains 168 and RO-NN-1 have an average of one genetic variant approximately every 50 bp, which allows identification of parental genomic contributions at a similar resolution. Small recombination events, particularly for closely-related strains such as these, may not result in any genetic change and will go undetected. Therefore, these measurements provide a lower bound on the recombination rate. Similarly, we chose to calculate the minimum insert size, based on the shortest length between recombined variants (Supplementary Figure S2). The true length of exchanged DNA may, in some cases, be several-fold longer. Finally, we cannot rule out that some genetic changes may be the result of more complicated DNA exchanges. A long insert may actually result from two adjacent recombination events, and it is difficult to precisely confirm the directionality of frequent fine-scale recombination events. In all cases, we have chosen the most parsimonious explanation.

### Identifying effects of genomic features on recombination

We next sought to identify genomic properties that might have influenced recombination. We hypothesized that differential methylation patterns in the two parents might bias recombination directionality and localization. Methylation analysis of our PacBio sequencing data confirmed the known GAGG<u>A</u>C methylation motif in strain 168 (39) and identified a different motif, A<u>A</u>GNNNNNNCR<u>T</u>C in RO-NN-1. The methyltransferase in strain 168 is not associated with a cognate restriction enzyme, and methylation in this strain is instead thought to influence transcriptional regulation (39). Conversely, strain RO-NN-1 encodes both a putative type I restriction enzyme targeting unmethylated DNA and a putative type IV restriction enzyme targeting methylated DNA. We therefore hypothesize that biased inheritance of RO-NN-1 genomic DNA in recombinant progeny is due to asymmetric enzymatic cleavage of the 168 chromosome in fused protoplasts.

To investigate more subtle influences on recombination, we examined the correlation between local recombination frequencies and several features of the genomic context, including proximity to methylation sites, local GC content, and SNP density (Figure 3). We performed these analyses using all recombination regions in the 168 HK x RO-NN-1 ME prototrophic progeny where we identified extensive genetic variability (Figure 1B). First, we hypothesized that double-strand breaks caused by DNA restriction might trigger increased homologous recombination near the restriction site. To test this hypothesis, we calculated distances between the boundaries of the recombination segments and the nearest methylation sites. Comparison of experimental data to the equivalent measurement for randomly permuted recombination regions showed no significant differences (Figure 3A). Thus, while methylation landscapes of the parental strains seem to determine directionality of recombination, they do not appear to affect the location of recombination events around the chromosomes of the offspring.

**Figure 3:**
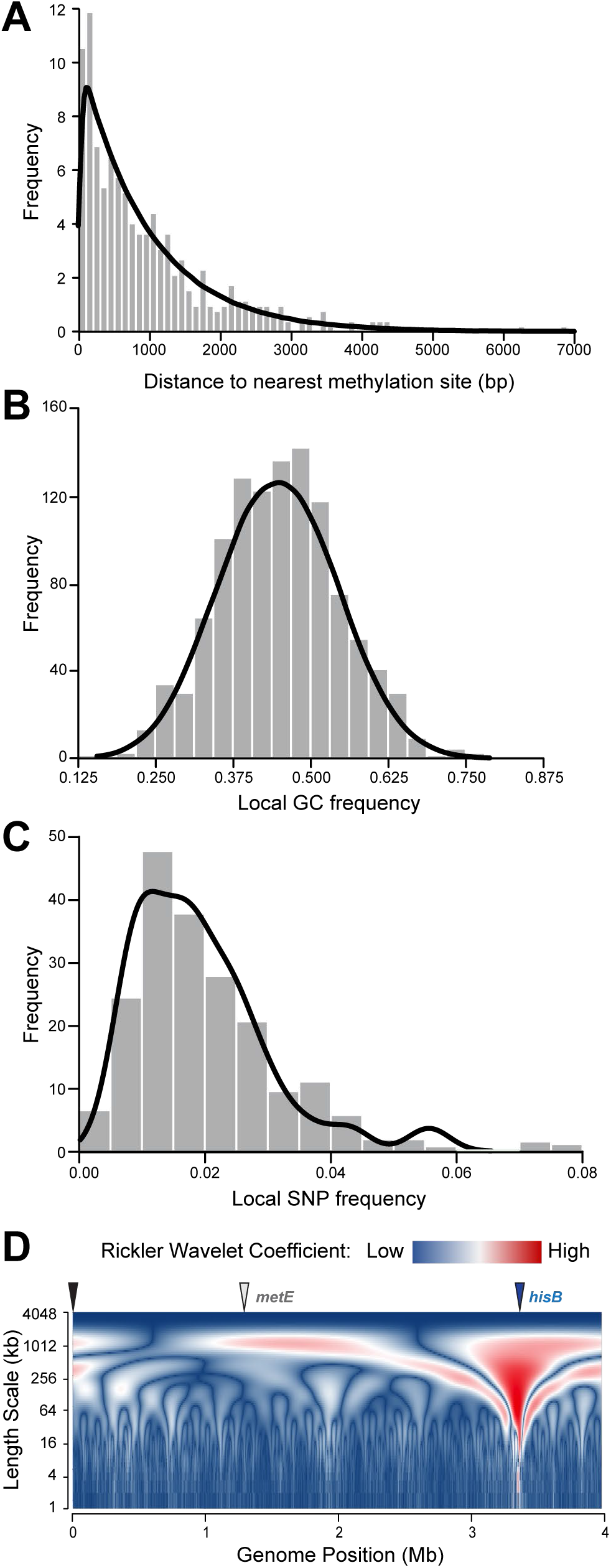
Genomic features do not affect recombination. (A-C) Genome properties were calculated for the complete set of 560 recombination sites in 168 HK x RO-NN-1 ME prototrophic progeny (grey histograms) and equivalent randomly permuted recombination sites (black lines). Features analyzed are (A) distance between the boundary of a recombination site and the nearest methylation site, (B) GC frequency in a 256 bp window spanning the recombination boundary, and (C) SNP frequency in the same 256 bp window. Differences between actual and permuted distributions were not significant. (D) Population-level recombination was analyzed across the genome using a Continuous Wavelet Transformation analysis with Ricker Wavelets. The wavelet coefficient is plotted for each combination of genomic position and length scale. High wavelet coefficients indicate deviations from the baseline at a particular combination of position and length scale. Genomic positions of the selection markers are indicated; this population selected for recombination at the *hisB* marker and against recombination at the metE marker. Only the recombination hotspot at *hisB* is evident.

In eukaryotes, a relationship between high GC content and elevated frequencies of recombination during meiosis has been well documented (40, 41). One possible explanation for this phenomenon is that regions rich in GC are more susceptible to formation of double strand breaks (41). We analysed GC richness of the homologous regions surrounding the recombination segments in the shuffled genomes and compared these results to GC content of regions flanking randomly-permuted recombination sites. We did not reveal any bias in recombination frequency towards GC content for the tested window sizes (2-512 bp, increasing 2^n^, Figure 3B shows a 256 bp window) (Figure 3B). These findings suggested that, at a fine scale, this aspect of the genomic environment might not be relevant for distribution of recombination events across the chromosome.

To evaluate the correlation between genetic diversity and recombination rates, we next estimated local SNP densities in regions flanking the recombination segments, in a similar fashion to the analysis of GC content. Rates of transformation by homologous recombination decrease exponentially as a function of sequence divergence, and efficient recombination is typically expected to require long homologous regions (42–44). As a result, we hypothesized that recombination would occur more frequently in regions of high local sequence identity. However, the local SNP frequency near recombination junctions did not differ significantly from random chromosomal locations in the tested window sizes (between 2 and 512 bp, 256 bp shown in Figure 3C). Similar to GC content, local SNP frequency does not appear to cause a significant bias in recombination patterns.

Patterns of recombination have been extensively investigated in eukaryotes. Distribution of meiotic recombination events across eukaryotic chromosomes is nonrandom. It has become clear that recombination predominantly occurs in specific regions of the genome known as recombination hotspots (45, 46). Since the genomic position and relevant length scale of a potential recombination hotspot is not known a priori, we used Continuous Wavelet Transformation analysis to simultaneously analyze average recombination frequencies of all potential positions and lengths (Figure 3D). We identified the known hotspot at the selection marker from the recessive parent (*hisB*) but could not detect any other biases. Wavelet analysis could also potentially detect regions with lower recombination frequencies than expected by chance. One known coldspot is located at the other selection marker (*metE*), but this location was not identified in our analysis. A higher average recombination frequency would be necessary to accurately detect coldspots.

Taken together, our analyses of the impact of genomic context on recombination rates did not reveal positive or negative associations. Other factors might also affect distribution of crossover events across the genome. Bacterial chromosomal DNA is organized into a compact structure called nucleoid by the cooperative action of DNA supercoiling and nucleoid associated proteins (47). Several lines of evidence suggested that rates of recombination could be affected by chromosome architecture. For example, analysis of site-specific recombination between regions scattered over the chromosome in *E. coli* demonstrated that intra-molecular recombination between different nucleoid macrodomains is highly restricted (48). Thus, some regions of the parental genomes might be randomly and temporarily more accessible for recombination in protoplast fusants which could explain the higher frequencies of local crossover events. Future investigation of the effect of DNA topology on recombination using the computational methods developed in this study might provide a mechanistic insight into genome-wide recombination patterns in bacteria.

### Effects of genetic distance on recombination

Strains 168 and RO-NN-1 are in the same subspecies and have approximately 98% average nucleotide identity (ANI) in shared genes (49). Genetic distance is an important physical factor that affects rates of recombination. To better understand the role of nucleotide identity on recombination parameters, we shuffled RO-NN-1 Δ*hisB*::*kan* Δ*metE*::*erm* with wild-type strains *B. subtilis* subsp. *subtilis* NCIB3610 (the parent of strain 168, with 98% ANI), *B. subtilis* subsp. *spizizenii* TU-B-10 (93% ANI) and *B. mojavensis* RO-H-1 (87% ANI). We were unable to generate recombinants using *B. velezensis* FZB42 (78% ANI). In each successful example, we isolated and resequenced approximately 16 strains from both potential recombinant genotypes, either Δ*hisB*::*kan metE*^+^ or Δ*metE*::*erm hisB*^+^. Previous analyses of single loci detected a log-linear relationship between genetic distance and recombination rates in *B. subtilis*, reaching saturation at about 8% sequence divergence between donor and recipient (43). Furthermore, the size of recombined regions decreased with increasing genetic distance (50). Surprisingly, in our study we observed no significant changes in the number of recombination events per strain (Figure 4A, Supplementary Figure S3 and Supplementary Table S1) or the size distribution of those recombined segments (Figure 4B). These results suggested that protoplast fusion might eliminate biases in recombination associated with transformation, for example by saturating the mismatch repair pathways.

**Figure 4:**
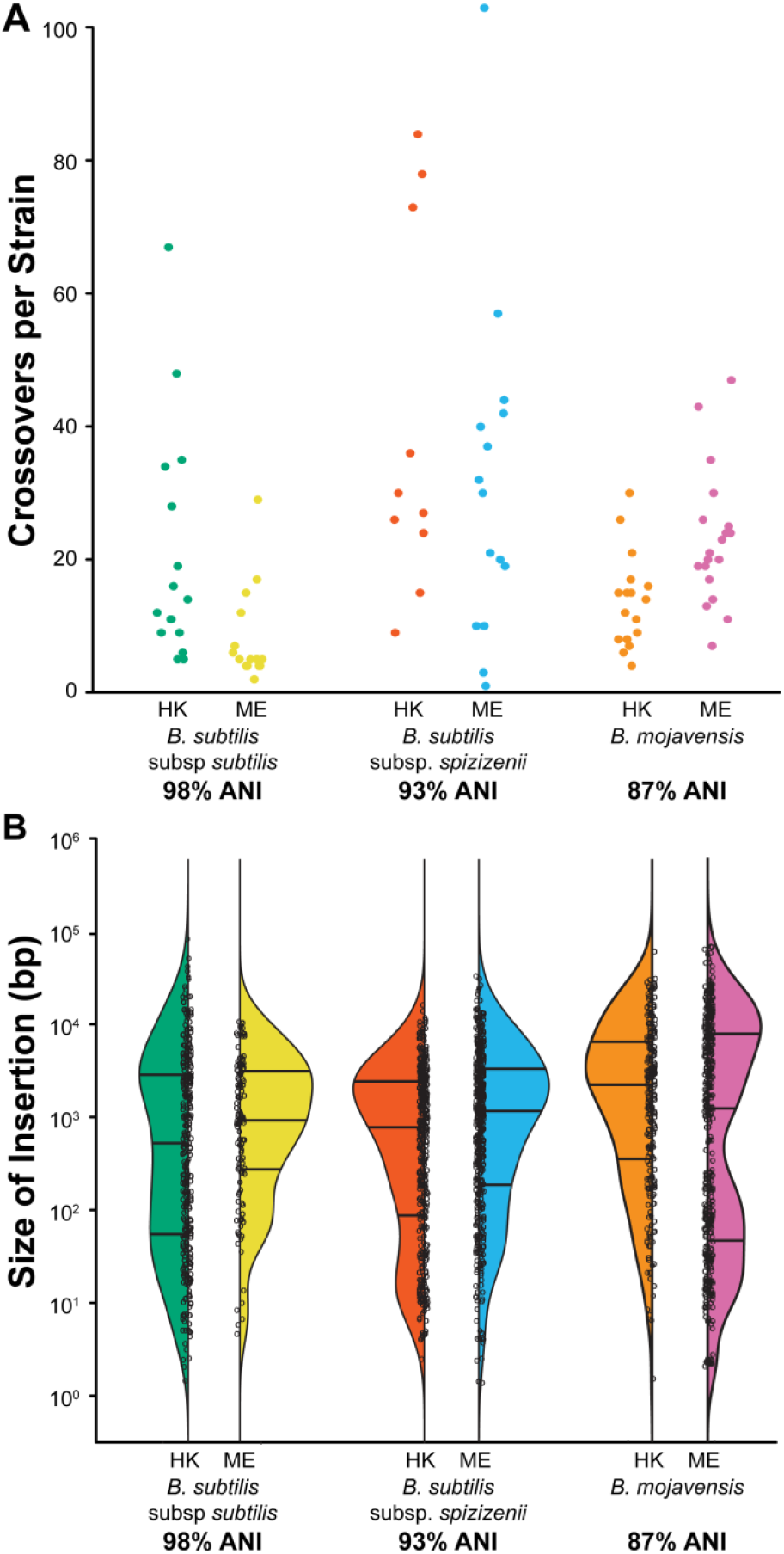
Protoplast fusion yields efficient homologous recombination across species boundaries. A double-resistant mutant of RO-NN-1 was crossed with prototrophic strains of varying genetic distance. No significant differences were observed in (A) the number of recombination events per strain or (B) the distribution of recombination event sizes. Horizontal lines in the violin plots show the median and interquartile range for each distribution.

In this work, we investigated the genomic consequences of protoplast fusion between *Bacillus* strains. We observed substantial un-selected recombination throughout the genome, for a broad range of fragment sizes. Restriction-modification systems strongly affected the directionality of transfer, but no other factors were identified that biased the local position of recombination events. While we were unable to obtain recombinants between strains with low levels of sequence identity, recombination was otherwise largely unaffected by variation in sequence identity between parental strains, even among strains classified as different species. When combined with the computational tools developed in this work, protoplast fusion provides a tractable method for studying homologous recombination at scale with minimal selection bias.

## AVAILABILITY

Source code for calculating subpopulation figures is available at GitHub: https://github.com/jstreich/Bsubstilis_QTL_Project_Aug2020

## ACKNOWLEDGEMENT

*Bacillus* strains were provided by Daniel Zeigler and the Bacillus Genetic Stock Center, through contributions from the initial depositors.

