## Supplementary information for "Protoplast fusion in *Bacillus* species produces frequent, unbiased, genome-wide homologous recombination"

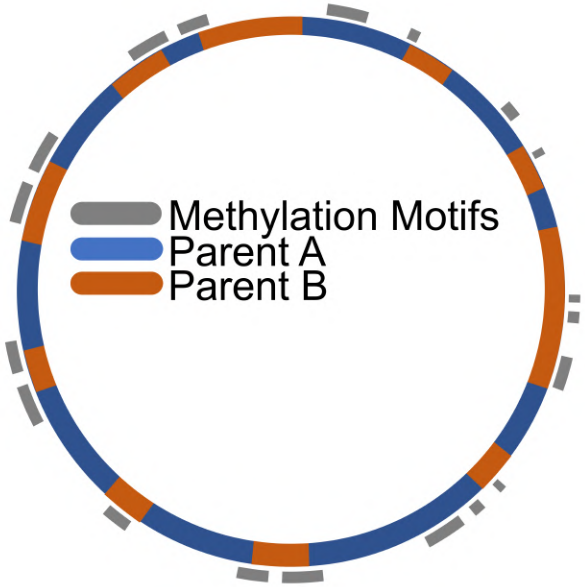


#### Actual Insertions

| 5′ Position | Insertion Size (3′ – 5′) | 3′ Position |
|---|---|---|
| 507,842 | 15,804 | 523,646 |
| 995,241 | 298 | 995,539 |
| … | … | … |
| 3,957,415 | 5,551 | 3,962,966 |

*n* = number of insertions

Randomization

#### *In-silico* Derived Insertions

| 5′ Positions Randomly drawn numbers | Insertion Size Actual Insertion Sizes Randomized Across Column | 3′ Position Random 5′ + laterally drawn insertion sizes |
|---|---|---|
| 12,844 | 5,551 | 18,395 |
| 1,858,488 | 298 | 1,858,786 |
| … | … | … |
| 3,915,020 | 15,804 | 3,930,824 |

*n* = number of insertions
1,000 iterations

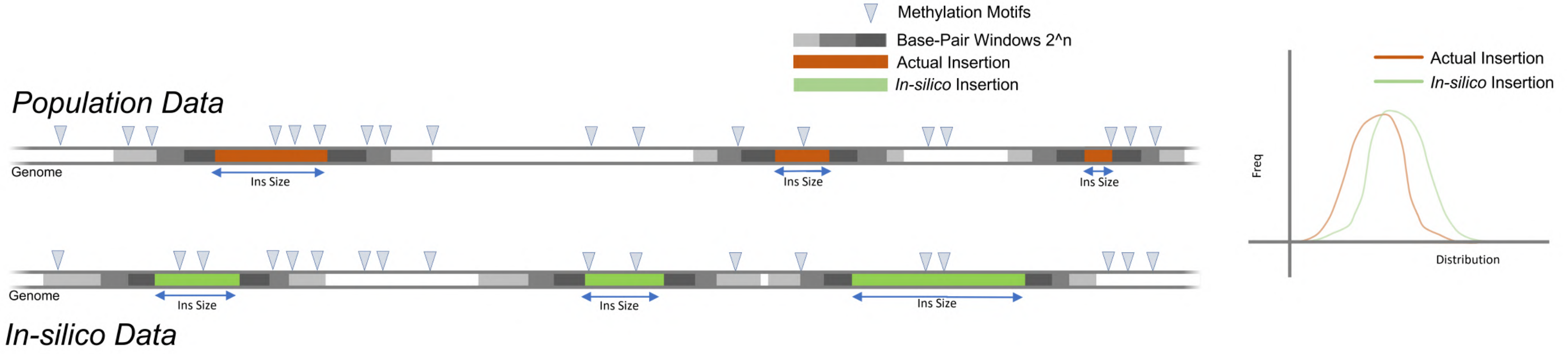


**Figure S1 Overview of workflow for evaluating association between recombination and local methylation patterns.** Actual inserts were called based on strings of continuous non-reference variants. The start and stop positions of these fragments were extracted from ped/map files and used for calculation of lengths of recombined regions. To generate *in-silico* fragments, random positions were drawn for each permutation iteration and randomly drawn sizes of actual recombined regions were added to random 5' positions creating 3' positions of comparable lengths. Exponentially increasing scans for number of motifs were used to compare proximity to methylation between boundaries of actual recombination regions and *in-silico* generated fragments.

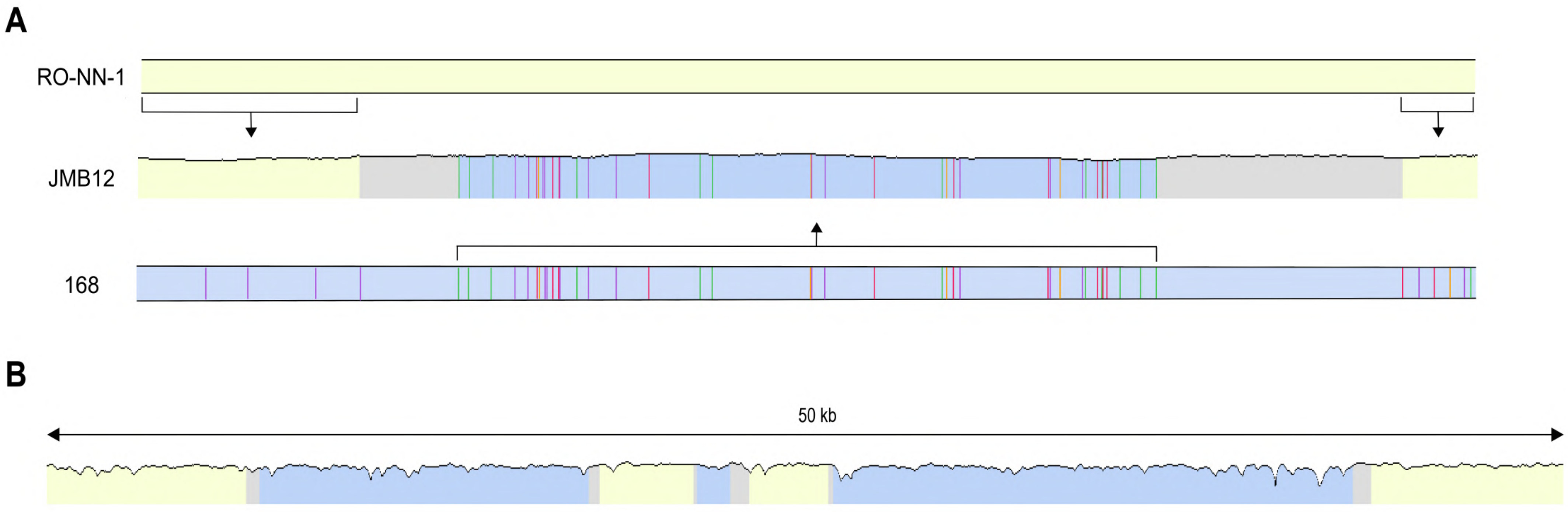


**Figure S2 Recombination events resulting from genome shuffling between *Bacillus subtilis* RO-NN-1 and 168.** (A) Assignment of recombination segment boundaries. Sequence reads of the recombinant strains were mapped to the reference genome of the dominant parent *B. subtilis* RO-NN-1 and single nucleotide polymorphisms (SNPs) inherited from the second parent *B. subtilis* 168 were identified. Regions encompassing DNA sequences between at least two consecutive 168-derived SNPs were defined as recombination regions. The reference genome sequences of strains RO-NN-1 and 168 are indicated as yellow and blue rectangles, respectively. The recombinant genome of strain JMB12 is presented as a sequence coverage histogram where RO-NN-1 and 168-derived regions are highlighted in yellow and blue. Segments of DNA flanking the recombination regions where no discriminatory SNPs between strains RO-NN-1 and 168 were identified are indicated in grey. Colored bars represent four different bases (A, red; T, green; C, purple; G, orange). (B) Recombination segments were clustered in small regions of the genome of each recombinant strain (see also Fig. 1B-D).

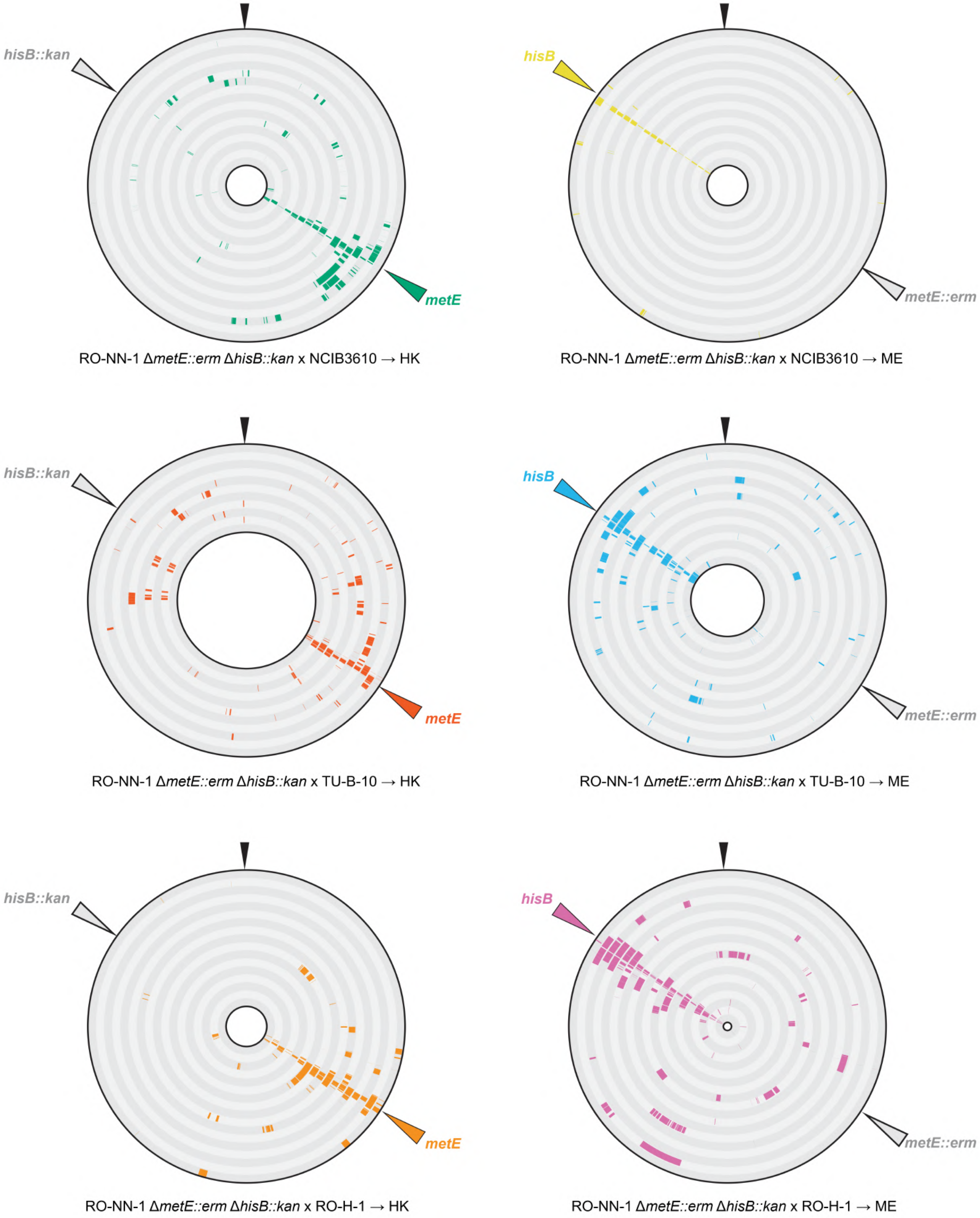


**Figure S3 Effect of genetic distance on genome-wide recombination patterns.** *B. subtilis* RO-NN-1 Δ*hisB*::*kan* ΔmetE::*erm* strain was crossed with wild-type *B. subtilis* subsp. *subtilis* NCIB3610 (98% ANI), *B. subtilis* subsp. *spizizenii* TU-B-10 (93% ANI) and *B. mojavensis* RO-H-1 (87% ANI). Histidine (HK) and methionine (ME) auxotrophic progeny were selected. Concentric circles represent resequenced individuals from each cross. The colored bars indicate recombined regions originating from NCIB3610, TU-B-10 and RO-H-1, with the remaining of the genome sequences coming from RO-NN-1. Green, yellow, red, blue, orange, and pink arrows show location of the selection markers. Black arrows indicate origin of replication.

### Table S1 Summary of recombination parameters in genome shuffled populations.

| Population | Number of Progeny | Total number of recombination events in population | Mean number of recombination events per strain | Mean size of recombined segment (bp) | Median size of recombined segment (bp) | Standard deviation of recombined segment size (bp) | % replaced genome (average) | Number of recombination event variance in population | Shapiro-Wilks test for number of recombination events p-value | Shapiro-Wilks test for number of recombination events *W* | Size of recombined segment variance in population | Shapiro-Wilks test for size of recombined segment p-value | Shapiro-Wilks test for size of recombined segment *W* |
|---|---|---|---|---|---|---|---|---|---|---|---|---|---|
| 168ME x RO-NN-1HK → WT | 18 | 560 | 31.1 | 5,847 | 1,092 | 11,111 | 4.5360 | 442.8187 | 0.0074 | 0.8525 | 123444865 | $<2.2\times10^{-16}$ | 0.5635 |
| 168ME x RO-NN-1HK → DR | 18 | 106 | 5.9 | 4,265 | 3,287 | 4,661 | 0.6260 | 103.1462 | $1.01\times10^{-6}$ | 0.5337 | 21723244 | $<2.2\times10^{-16}$ | 0.8180 |
| 168HK x RO-NN-1ME → WT* | 18 | - | - | - | - | - | - | - | - | - | - | - | - |
| 168HK x RO-NN-1ME → DR | 18 | 600 | 33.3 | 13,027 | 8,639 | 14,699 | 2.9960 | 52.06536 | 0.0060 | 0.8405 | 216064902 | $4.39\times10^{-13}$ | 0.4057 |
| RO-NN-1HKME x NCIB3610 → HK | 15 | 472 | 31.5 | 242 | 1,113 | 3,336 | 1.5830 | 759.8909 | 0.0596 | 0.8611 | 11131027 | $<2.2\times10^{-16}$ | 0.7194 |
| RO-NN-1HKME x NCIB3610 → ME | 15 | 469 | 31.3 | 3,415 | 1,502 | 5,320 | 2.6630 | 657.0667 | 0.0394 | 0.8746 | 28300406 | $<2.2\times10^{-16}$ | 0.6361 |
| RO-NN-1HKME x TU-B-10 → HK | 17 | 234 | 13.8 | 6,588 | 2,868 | 9,984 | 2.2620 | 49.19118 | 0.2390 | 0.9324 | 99674735 | $<2.2\times10^{-16}$ | 0.6616 |
| RO-NN-1HKME x TU-B-10 → ME | 18 | 438 | 24.3 | 7,450 | 1,604 | 12,879 | 4.5200 | 103.0526 | 0.1763 | 0.9304 | 165873711 | $<2.2\times10^{-16}$ | 0.6213 |
| RO-NN-1HKME x RO-H-1 → HK | 16 | 338 | 21.1 | 4,119 | 669 | 9,692 | 2.1710 | 297.8603 | 0.0019 | 0.7984 | 93944495 | $<2.2\times10^{-16}$ | 0.4386 |
| RO-NN-1HKME x RO-H-1 → ME | 16 | 131 | 8.2 | 2,705 | 1,194 | 3,273 | 0.5520 | 48.22059 | $1.46\times10^{-4}$ | 0.7088 | 10710508 | $<2.2\times10^{-16}$ | 0.7690 |

Columns 9-11 and 12-14 show statistical analyses of number of recombination events and recombined segment sizes. Shapiro-Wilk tests were performed to test distribution normality. Each population had non-normally distributed data.

* 168HK x RO-NN-1ME prototrophic population has been contaminated and not analyzed further (see text for details).

**Table S2 Comparative analyses of number of recombination events between 168ME x RO-NN-1HK WT, 168ME x RO-NN-1HK DR and 168HK x RO-NN-1ME DR shuffled populations (see Figure 2A).**

| Populations | Wilcoxon Test | | Kolmogorov-Smirnov Test | | T-Test | | F-Test | |
|---|---|---|---|---|---|---|---|---|
| | p-value | W | p-value | D | p-value | t* | p-value | F |
| 168ME x RO-NN-1HK WT compared to 168HK x RO-NN-1ME DR | 1 | 180 | 0.9718 | 0.1579 | 0.7560 | -0.29926 | 0.5960 | 0.8901 |
| 168ME x RO-NN-1HK WT compared to 168ME x RO-NN-1HK DR | $6.81 \times 10^{-6}$ | 332 | $1.44 \times 10^{-5}$ | 0.7895 | $1.412 \times 10^{-3}$ | 4.4576 | 0.0017 | 4.2931 |
| 168HK x RO-NN-1ME DR compared to 168ME x RO-NN-1HK DR | $8.98 \times 10^{-6}$ | 330 | $1.44 \times 10^{-5}$ | 0.7895 | $9.73 \times 10^{-5}$ | 4.6243 | $8.33 \times 10^{-3}$ | 4.8231 |

$p<0.001$ statistically significant
